# Stepwise maturation of the peptidyl transferase region of human mitoribosomes

**DOI:** 10.1101/2021.03.29.437532

**Authors:** Tea Lenarčič, Mateusz Jaskolowski, Marc Leibundgut, Alain Scaiola, Tanja Schönhut, Martin Saurer, Richard Lee, Oliver Rackham, Aleksandra Filipovska, Nenad Ban

**Affiliations:** Department of Biology, Institute of Molecular Biology and Biophysics, ETH Zurich, Otto-Stern-Weg 5, CH-8093 Zurich, Switzerland; Harry Perkins Institute of Medical Research, ARC Centre of Excellence in Synthetic Biology, QEII Medical Centre, University of Western Australia, Telethon Kids Institute, Northern Entrance, Perth Children’s Hospital, 15 Hospital Avenue, Nedlands, Western Australia 6009, Australia; Curtin Health Innovation Research Institute and Curtin Medical School, Curtin University, Bentley, Western Australia 6102, Australia

**Author notes:** Correspondence should be addressed to N.B. These authors contributed equally to this work (Tea Lenarčič, Mateusz Jaskolowski).

## Abstract

Mitochondrial ribosomes are specialized for the synthesis of membrane proteins responsible for oxidative phosphorylation. Mammalian mitoribosomes diverged considerably from the ancestral bacterial ribosomes and feature dramatically reduced ribosomal RNAs. Structural basis of the mammalian mitochondrial ribosome assembly is currently not understood. Here we present eight distinct assembly intermediates of the human large mitoribosomal subunit involving 7 assembly factors. We discover that NSUN4-MTERF4 dimer plays a critical role in the process by stabilizing the 16S rRNA in a conformation that exposes the functionally important regions of rRNA for modification by MRM2 methyltransferase and quality control interactions with a conserved mitochondrial GTPase MTG2 that contacts the sarcin ricin loop and the immature active site. The successive action of these factors leads to the formation of the peptidyl transferase active site of the mitoribosome and the folding of the surrounding rRNA regions responsible for interactions with tRNAs and the small ribosomal subunit.

---

Human mitochondrial ribosomes (mitoribosomes) are responsible for translation of 13 oxidative phosphorylation (OXPHOS) proteins, encoded in the mitochondrial genome^1,2^. Due to their unusual architectural features^3–5^ and the requirement to coordinate mitochondrial ribosomal RNA (rRNA) synthesis with import of all ribosomal proteins, their assembly is anticipated to involve mitochondrial-specific pathways and participation of both conserved and mitochondrial-specific maturation factors. Many factors, including conserved and essential GTPases and methyltransferases participate in human mitoribosome assembly^1^. They ensure production of translationally competent ribosomes and defects in the mitochondrial translation machinery lead to a range of severe human pathologies^6^. Although human mitoribosome assembly has been extensively investigated using a combination of biochemical and high-throughput approaches^1,7–9^, structural understanding of this process is currently limited to a late assembly intermediate with a bound MALSU1–L0R8F8–mt-ACP module^10^.

To better understand the structural basis of human mitoribosome maturation, we isolated large mitoribosomal subunit (mt-LSU) assembly intermediates and investigated their composition and structures using cryo-electron microscopy (cryo-EM). We determined structures of eight distinct states of the human mt-LSU where a total of 7 assembly factors were bound in different combinations. Our results reveal the conformational changes that allow successive modification and maturation of the functionally important regions of rRNA. The structural data supported by biochemical evidence provide an explanation for the role of the essential NSUN4-MTERF4 heterodimer in the process and specifically in the maturation of the functionally important, tRNA interacting, P loop through interplay with the conserved GTPase MTG2. The obtained results allow us to propose a stepwise maturation pathway of the functionally important regions in human mitochondrial large ribosomal subunit.

## Results

### Structure of assembly intermediates of the human mitoribosomal large subunit

To purify assembly intermediates of the human mitochondrial large ribosomal subunit (mt-LSU), we transfected human embryonic kidney cells with a tagged mitochondrial GTPase 1 (MTG1) (**Supplementary Fig. 1**), which is essential for production of functional mitoribosomes due to its involvement in the late stages of mt-LSU assembly prior to monosome formation^11,12^. The affinity-purified sample was investigated using single particle cryo-electron microscopy (cryo-EM) to reveal several mt-LSU-like structures containing the entire set of ribosomal proteins. Furthermore, density for several additional proteins was observed at the intersubunit side bound to the rRNA in an immature conformation. Using focused classification around those additional features, we were able to obtain high resolution reconstructions of 8 distinct cryo-EM classes corresponding to assembly intermediate states of the mt-LSU (**Supplementary Fig. 2 and 3**). Two of the classes resemble the structures of the assembly intermediates described previously, where a MALSU1–L0R8F8–mt-ACP module is bound at the intersubunit face, while the rRNA is either disordered or in a nearly mature state^10^. The remaining six classes correspond to novel assembly intermediates (**Supplementary Fig. 2**). We characterised three of these intermediates, resolved to 2.88 Å, 3.05 Å and 3.05 Å and referred to as states A, B and C, respectively (**Fig. 1 and Supplementary Table 1**), as the key steps in the late stages of mt-LSU maturation, during which functionally important regions of the rRNA progressively mature.

**Fig. 1.**
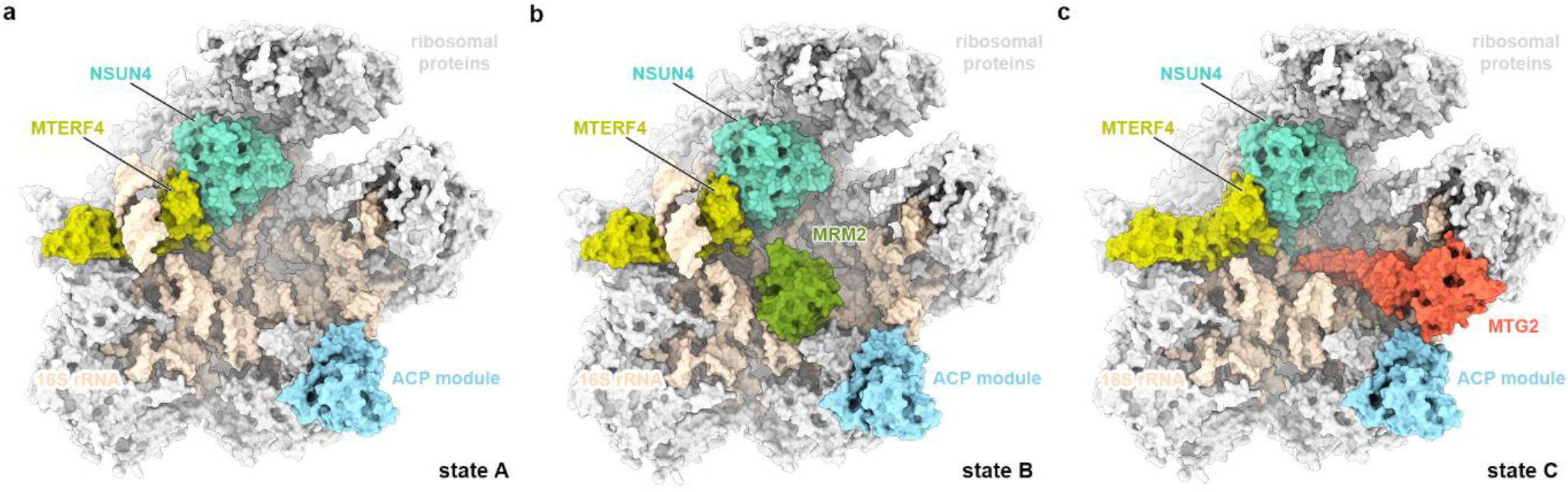
Structures of the key human mitoribosomal large subunit assembly intermediates. The molecular structures of states A (a), B (b) and C (c) are shown in surface representation from the intersubunit side. Ribosomal proteins are depicted in light grey, 16S rRNA in beige and assembly factors NSUN4, MTERF4, MRM2, MTG2 and the MALSU1–L0R8F8–mt-ACP module in teal, light green, green, red and blue, respectively.

### NSUN4-MTERF4 stabilizes rRNA in a conformation that exposes the active site region

In addition to the previously characterized MALSU1–L0R8F8–mt-ACP module^10^, state A contains at the intersubunit side, below the central protuberance (CP), a dimer of NSUN4 and MTERF4, which were both previously identified as mitoribosomal assembly factors^13,14^ (**Fig. 1a**). Although tagged MTG1 was used for affinity purification of ribosomal assembly intermediates, the factor was not sufficiently ordered on the mt-LSU to be structurally interpreted in any of the observed states. The NSUN4–MTERF4 dimer forms extensive interactions with rRNA and ribosomal proteins by contacting rRNA helices H66, H75, H81, H87 and H93, and keeps the C-termini of uL2m and mL48 in an immature conformation (**Fig. 2a**). Interestingly, NSUN4 is an RNA m^5^C methyltransferase that has been implicated in assembly of both small and large mitoribosomal subunits, however, biochemical data suggested that *in vivo* it only modifies the rRNA of the small subunit^14^. Consistently, no RNA substrate was found in the active site of NSUN4 where a clearly visible S-adenosyl-methionine (SAM) cofactor was bound (**Fig. 2b and Supplementary Fig. 4**). Since NSUN4 lacks an RNA substrate recognition domain that is present in bacterial homologues^13^, it was proposed that it relies on interactions with MTERF4 to be targeted to the mt-LSU to regulate ribosome maturation^13,14^, as we now observe in the mt-LSU bound state.

**Fig. 2.**
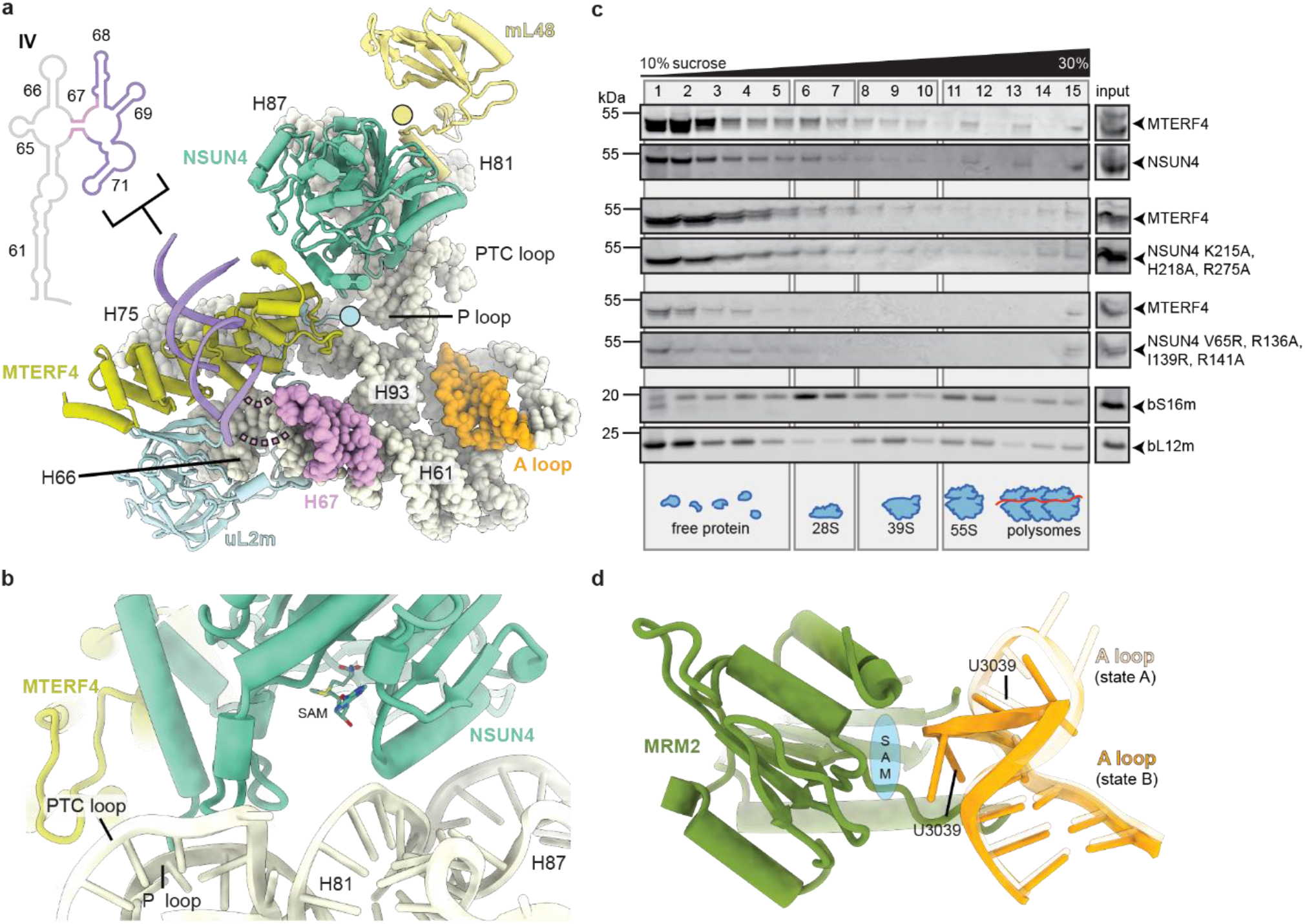
Interactions of NSUN4 and MTERF4 with the immature human mitoribosomal large subunit. (a) The NSUN4–MTERF4 heterodimer (teal and light green, respectively) forms a number of contacts with the mitochondrial large subunit *via* rRNA (beige spheres) and ribosomal proteins, resulting in rearrangements of the mL48 (yellow) and uL2m (light blue) C-termini (highlighted as circles), as well as stabilization of the H68-H71 region (violet) of the 16S rRNA domain IV. The connection from helix H67 (pink) to H68-H71 is shown as dashed lines. The A loop, which is modified by MRM2 in state B, is colored orange. (b) The NSUN4 (teal) active site with bound S-adenosyl-methionine (SAM) cofactor is shown together with nearby rRNA helices H81, H87, P loop and PTC loop (beige cartoon). MTERF4 is also shown for orientation (light green). (c) Effects of mutations in NSUN4 or MTERF4 on their association with mitoribosomal fractions. A continuous 10%–30% sucrose gradient was used to separate mitochondrial lysates from HEK 293T cells expressing MTERF4 and either wild-type or mutated version of NSUN4 to determine their distribution and co-migration with mitochondrial ribosomal fractions. The small and large ribosomal subunit and polysomes in mitochondria isolated from wild-type cells were followed by immunoblotting for mitochondrial ribosomal protein markers of the small (bS16m) and large (bL12m) ribosomal subunits. The input, mitochondrial lysate, was used as a positive control. (d) Rearrangement of the A loop (light orange: state A; orange: state B) upon the MRM2 methylation event. The likely position of MRM2 SAM cofactor, which is not visible in our structure, is schematically shown in blue.

The MTERF4 protein folds into a bent α-solenoid that binds with its convex region to the surface of the immature subunit and exposes its positively charged concave region towards the outside, where we observe a segment of double helical RNA bound (**Supplementary Fig. 5**). Although we could not assign the identity of RNA nucleotides in this region, continuous density can be traced from rRNA helix H67, suggesting that the additional density belongs to the H68-H71 region of the rRNA (**Fig. 2a and Supplementary Fig. 5**). In the mature mt-LSU these helices form the front of the peptidyl transferase active site cleft. The interactions with MTERF4 expose the immature rRNA regions corresponding to the peptidyl transferase (PTC) loop that forms the active site of the ribosome, as well as the A (H92) and P (H80) loops responsible for binding the acceptor end of tRNAs and positioning substrates for the peptidyl transferase reaction in the active site of the mature mt-LSU.

To better understand the role of both MTERF4 and NSUN4 in the context of the mt-LSU assembly, we designed a series of mutants based on our structural results with deletions in key interaction regions. The mutant proteins were then investigated with respect to their ability to form dimers and associate with the mitoribosome using continuous sucrose gradients in cells expressing a FLAG-tagged MTERF4 and HA-tagged NSUN4. Wild-type MTERF4 and NSUN4 associate with each other and co-migrate with the large subunit of the mitoribosome, confirming previous findings^13–16^. Two different mutants were designed to disrupt NSUN4 interactions with the 16S rRNA or MTERF4. The first NSUN4 mutant, bearing a triple mutation K215A, H218A and R275A located at the interface between the rRNA and NSUN4 (**Supplementary Fig. 5**), was designed to test the contribution of NSUN4 to binding of the complex to the 16S rRNA of the immature mt-LSU. This mutation neither reduced the association of NSUN4 with MTERF4 nor their co-migration with the large subunit (**Fig. 2c**), indicating that MTERF4 plays a predominant role in delivering the complex to the immature subunit. A second NSUN4 mutant carrying a quadruple mutation of residues V65R, R136A, I139R and R141A (**Supplementary Fig. 5**), designed to break the dimer between NSUN4 and MTERF4, completely abolished their interaction as shown previously^16^, and also prevented binding of either of the two proteins to the large subunit (**Fig. 2c**). We conclude that formation of a stable NSUN4–MTERF4 heterodimer is critical for their function in mitoribosome assembly.

### MRM2 methylates the U3039 in the A loop before it adopts mature conformation

While the above described state A contains only the MALSU1–L0R8F8–mt-ACP module and NSUN4–MTERF4 dimer, in state B we additionally observe the MRM2 methyltransferase bound to the rRNA in an optimal position for methylation of its target nucleotide U3039 within the A loop of the 16S rRNA^17,18^ (**Fig. 1b**). The A loop is repositioned such that the 2’-O-ribose of nucleotide U3039 faces the active site of MRM2, although we do not observe density for a SAM cofactor (**Fig. 2d**). The binding site for the MRM2 methyltransferase, as observed in state B, is occupied by helix H71 in the mature mt-LSU (**Supplementary Fig. 6**). This implies a temporal order of maturation events where U3039 methylation must take place before helix H71 assumes its mature conformation.

### MTG2 and NSUN4 interact with the P loop in a tweezer-like manner

In state C (**Fig. 1c**), MRM2 is replaced by MTG2, a GTPase conserved from bacteria to eukaryotes. The bacterial homolog (ObgE) has been recently visualized on the native assembly intermediates of the bacterial large subunit^19^. MTG2 was proposed to play a key role in the human mt-LSU assembly as a final quality control checkpoint protein^20^. Furthermore, biochemical experiments showed that it interacts with the mt-LSU at the same time as MRM2, MTERF4, MTG1 and MALSU1^20^. Our structure reveals that, in the presence of NSUN4, MTERF4 and the MALSU1–L0R8F8–mt-ACP module, MTG2 binds to the mt-LSU in a position, from which it can simultaneously check two key regions of the large ribosomal subunit: the GTPase associated center (GAC) that plays a key role in stimulating GTP hydrolysis of translational GTPases and the catalytic peptidyl transferase center (PTC) (**Fig. 3a**).

**Fig. 3.**
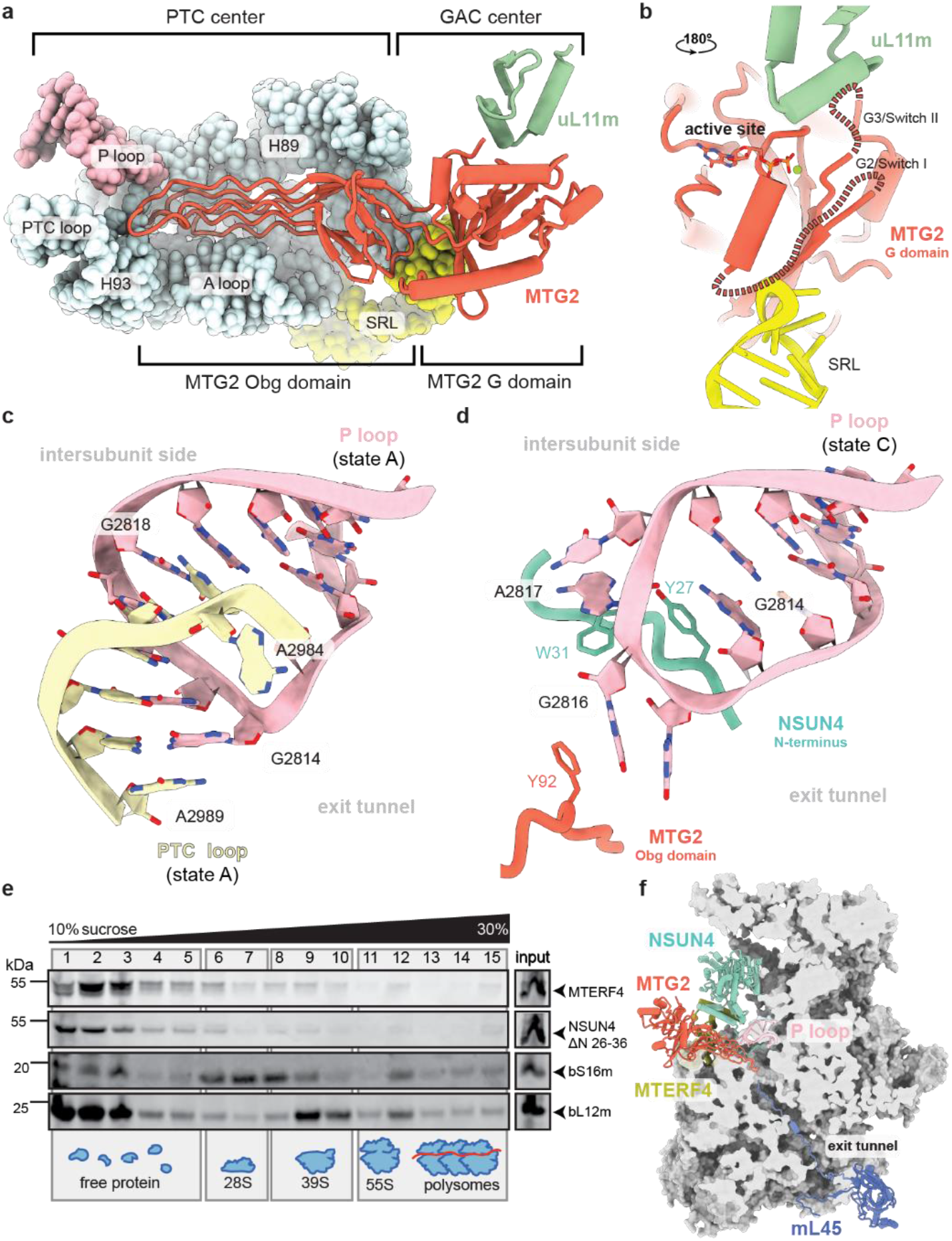
Mitochondrial GTPase MTG2 interacts with the functional regions of the immature human mitoribosomal large subunit. (a) Interaction of the MTG2 (red cartoon) with the 16S rRNA and ribosomal proteins in state C. The N-terminal Obg domain of MTG2 contacts the peptidyl transferase center (PTC) region of the 16S rRNA (light cyan, PTC helices depicted individually, P loop in pink), whereas the G domain associates with the ribosomal GTPase associated center (GAC) components uL11m (green) and the sarcin-ricin loop (SRL). (b) Detailed view of G domain interactions with the GAC rotated by 180° relative to panel (a). The Switch loops II and III are schematically indicated as dashed lines. The color key is the same as in panel (a). (c) The 16S rRNA PTC loop (light yellow) stacking with the P loop (pink) as a result of an immature rRNA arrangement in state A. (d) Specific interactions of the NSUN4 N-terminal tail (teal) and the MTG2 Obg domain (red) with the P loop (pink). Amino acid residues involved in coordinating the P loop are highlighted. (e) Effects of NSUN4 ΔN 26-36 mutant on the association with mitoribosomal fractions and 55S monosome formation. A continuous 10%–30% sucrose gradient was used to separate mitochondrial lysates from HEK 293T cells expressing the MTERF4 and NSUN4 mutant to determine their distribution and co-migration with mitochondrial ribosomal fractions. The small and large ribosomal subunit and polysomes in mitochondria isolated from NSUN4 ΔN 26-36 transfected cells were followed by immunoblotting for mitochondrial ribosomal protein markers of the small (bS16m) and large (bL12m) ribosomal subunits. The input, mitochondrial lysate, was used as a positive control. (f) Cross-section of the mitochondrial large subunit state C assembly intermediate. Spatial arrangement of assembly factors NSUN4 (teal), MTERF4 (light green) and MTG2 (red) ensure probing the P loop (pink) and the entrance to the nascent polypeptide tunnel in the mitochondrial large subunit assembly intermediate (gray). The mL45 (blue) N-terminal tail occupies the exit tunnel, contributing to an inactive state of the subunit.

On the side of the GAC, the conserved Ras-like G domain of MTG2 interacts with sarcin-ricin loop (SRL) H95 and ribosomal protein uL11m (**Fig. 3b**). The G domain is oriented such that the active site, including the only partially resolved G2/Switch I and G3/Switch II loops involved in GTP binding and hydrolysis, faces towards the SRL and uL11m. Such positioning suggests that the G domain of MTG2 would be able to monitor the correct conformation of the GAC region in the final stages of mt-LSU maturation.

On the other side, the N-terminal Obg domain of MTG2 stabilises the mature conformation of the PTC-forming helices H81, H89, H90 as well as the A and PTC loops and indirectly H71, which are in immature conformations in states A and B (**Supplementary Fig. 7**). The most prominent rRNA rearrangement involves the PTC loop, which in state A surprisingly forms a helix with the P loop of the 16S rRNA (**Fig. 3c**). As the PTC loop matures in state C, it withdraws from the P loop, which is now held in immature conformation by contacts with assembly factors MTG2 and NSUN4 (**Fig. 3d**). These interactions encompass the Obg domain of MTG2 and the N-terminal tail of NSUN4, which is disordered in the absence of MTG2. The two factors grip the immature P loop from two sides in a tweezer-like manner involving aromatic residues Tyr27 and Trp31 of NSUN4 that stack with the P loop nucleotides G2814 and A2817, whereas MTG2 contributes Phe92 to interact with the P loop nucleotide G2816 (**Fig. 3d**). While deletion of residues 26-36 at the N-terminal region of NSUN4 did not reduce the association of NSUN4 with MTERF4 or mt-LSU, it reduced the levels of mature mitoribosomes in a dominant negative manner (**Fig. 3e**) compared to the control (**Fig. 2c**). Taken together, our structural and biochemical results reveal direct contribution of NSUN4 N-terminal tail to the maturation of the mt-LSU.

The above-mentioned P loop is the sole element of the 16S rRNA domain V that remains immature in state C and, together with helices H68 and H69 of domain IV that are still bound to MTERF4, the only area in the mt-LSU rRNA that still needs to mature (**Supplementary Fig. 7**). Interestingly, besides keeping the P loop in a distinct immature conformation, MTG2 also samples the entrance to the nascent polypeptide tunnel with one of its Obg domain loops (**Fig. 3f**). At exit side of the ribosomal tunnel we observe that mitoribosomal protein mL45 inserts its N-terminal tail into the tunnel, as observed for the non-translating mature mitoribosomes^2,21^, and reaches almost to the Obg domain of MTG2. Consequently, the two proteins sample virtually the entire length of the exit tunnel and may play a role in facilitating proper folding of proteins and rRNA elements forming the mitoribosomal nascent polypetide tunnel during mt-LSU maturation.

### Stepwise maturation of the human mitochondrial large subunit

Visualisation of 8 distinct assembly intermediates of the human mt-LSU that reveal the interdependence of assembly factors and the role of the rRNA in the process allows us to propose a model for stepwise maturation of the ribosomal active site (**Fig. 4 and Supplementary Fig. 8**). Binding of the NSUN4–MTERF4 dimer to the immature, but compositionally complete mt-LSU, sequesters the flexibly disposed H68-H71 region to expose the active site and allow access of factors that modify the rRNA and check its conformation. This conformation is recognised by the methyltransferase MRM2 that modifies nucleotide U3039 in the A loop of the rRNA. After the methylation of U3039, MRM2 dissociates and the quality checkpoint GTPase MTG2 binds to the P loop in an NSUN4-dependent manner to facilitate maturation of the PTC and to check the functionality of the GAC of the ribosome. Once the rRNA assumes its native or nearly native conformation, the assembly factors dissociate and a mature, translationally competent mt-LSU is formed.

**Fig. 4.**
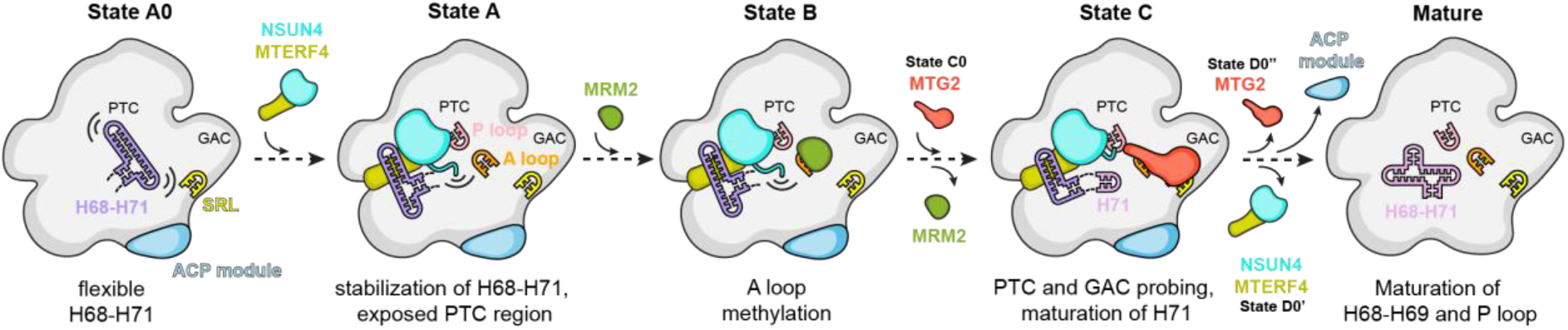
Model for stepwise maturation of the human mitoribosomal large subunit aided by assembly factors. Eight classes, corresponding to distinct assembly states in this study, allow us to propose the order of events in the late stages of the human mitoribosomal large subunit assembly. Sequestering of the H68-H71 16S rRNA by assembly factors NSUN4 and MTERF4 exposes the functionally important regions of the large subunit that allows MRM2 to modify its target nucleotide in the A loop. Dissociation of MRM2 is followed by association of MTG2, a mitochondrial GTPase that performs a final quality check of the peptidyl transferase center (PTC) region as well as the sarcin-ricin loop (SRL) in the GTPase associated center (GAC). Dissociation of all assembly factors results in completion of the rRNA maturation and formation of a translationally competent particle.

### Concluding remarks

Our data provides the basis for understanding the late stages of human mt-LSU maturation at a structural level and reveals the key role of the NSUN4 and MTERF4 to induce a conformation of the subunit ready for subsequent modification and maturation steps. These results complement the discovery of a remarkably complex mitoribosomal assembly machinery in trypanosomal mitochondria^22– 25^ and suggest that the divergent mitochondrial ribosomes are likely to involve equally diverse set of assembly factors across different species. These results are also biomedically important as defects in the mitochondrial translational apparatus cause a decline in mitochondrial function that is associated with development of neurodegenerative diseases^26^ and diverse progressive and fatal genetic disorders^27,28^.

## Methods

### Transient expression of MTG1-3xFLAG in HEK 293 EBNA cells

HEK 293 EBNA embryonic kidney cells, adapted to suspension growth in serum-free Ex-Cell medium, were obtained from the protein production and structure core facility at EPFL. The cell line was cultured at 37 °C under 4.5 % CO_2_ in EX-CELL® 293 Serum-Free Medium for HEK 293 Cells (Sigma), supplemented with 4 mM L-glutamine. The cells were not tested for mycoplasma contamination.

The pcDNA3.1(+) plasmid encoding the C-terminally 3xFLAG-tagged MTG1 (UniProt ID Q9BT17) was ordered from GenScript. The inserted sequence was verified using a CMV forward primer at Microsynth. Cells with a concentration of 10^6^ cells/mL were transfected with 1.5 mg of DNA per liter of the culture using transfection reagent 40 kDa PEI MAX (Polysciences, Inc.) in a 1:2 ratio. Cells were harvested and mitochondria were isolated 72 h post transfection.

### Isolation of mitochondria from HEK 293 EBNA cells

Mitochondria were isolated as previously described^1,2^ with few modifications. Briefly, the cell pellets were resuspended in ice-cold RSB hypo buffer (10 mM Tris-HCl pH 7.5, 10 mM NaCl and 1.5 mM MgCl_2_) and allowed to swell for 10 min. The swollen cells were opened with several strokes of a dounce homogenizer followed by immediate addition of 2.5x MS buffer (12.5 mM Tris-HCl pH 7.5, 525 mM mannitol, 175 mM sucrose, 2.5 mM EDTA and 2.5 mM DTT) to a final concentration of 1x MS buffer (5 mM Tris-HCl pH 7.5, 210 mM mannitol, 70 mM sucrose, 1 mM EDTA and 1 mM DTT). The homogenate was clarified at 1,300 g and 4 °C for 10 min, followed by another 10 min centrifugation at 7,500 g and 4 °C. Finally, the supernatant and the crude mitochondria fraction were separated after centrifugation at 9500 g and 4 °C for 10 min. The mitochondria pellet was resuspended in 20 mM HEPES-KOH pH 7.6, 250 mM sucrose and 1 mM EDTA and applied to a layered sucrose gradient, consisting of 15 %, 23 %, 32 % and 60 % (w/v) sucrose solutions in 20 mM HEPES-KOH pH 7.6 and 1 mM EDTA. After 70 min ultracentrifugation at 60,000 x g and 4 °C using SW 32 Ti rotor, mitochondria band between 32 % and 60 % (w/v) sucrose solution was carefully collected, flash frozen in liquid nitrogen and stored at -80 °C until use.

### Preparation of MTG1-3xFLAG tagged mitochondrial ribosomes

Upon thawing of the mitochondria, 1.5 volumes of lysis buffer (20 mM HEPES-KOH pH 7.6, 100 mM KCl, 20 mM MgCl_2_, 1.6 % Triton X-100, supplemented with 1x cOmplete EDTA-free protease inhibitor cocktail (Roche)) was added, and mitochondria were lysed using a dounce homogenizer. Membranes were further solubilized by stirring for 15 min at 4 °C. The lysate was clarified by centrifugation at 20,800 x g for 15 min at 4 °C. Next, the supernatant was incubated for 1.5-2 h at 4 °C with Anti-FLAG M2 affinity gel (Sigma) while gently mixing. The Anti-FLAG M2 affinity gel (Sigma) was pre-equilibrated with 3 sequential column volumes of 0.1 M glycine HCl pH 3.5 and washed with 10 volumes of TBS and wash buffer (20 mM HEPES-KOH pH 7.6, 100 mM KCl and 20 mM MgCl_2_). After collecting the flow-through, the beads were washed with 10 column volumes of wash buffer. Bound mitoribosomes were eluted 3 times using elution buffer (20 mM HEPES-KOH pH 7.6, 100 mM KCl, 20 mM MgCl_2_ and 100-200 µg/mL 3xFLAG peptide), each time preceded by a 10-15 minutes incubation at 4 °C with gentle mixing. Eluted fractions were pooled and subjected to a 2.5 h ultracentrifugation at 135,500 x g at 4 °C using a TLA-55 rotor (Beckman-Coulter). Finally, the mitoribosome pellet was resuspended in mito-resuspension buffer (20 mM HEPES-KOH pH 7.6, 100 mM KCl, 20 mM MgCl_2_ and 1 mM DTT), yielding mt-LSU at a concentration of approximately 40 nM. Samples from the key steps of the purification were subjected to western blot analysis (anti-3x-FLAG antibody A8592, Sigma) (**Supplementary Fig. 1**).

### Expression of NSUN4 variants in HEK 293T cells

Expression cassettes for NSUN4 variants were synthesized from overlapping oligonucleotides and cloned into pTwist CMV (Twist Bioscience). All NSUN4 variants were expressed as fusions to a C-terminal HA tag, while MTERF4 was C-terminally FLAG tagged.

Human embryonic kidney (HEK 293T) cells were cultured at 37 °C in humidified 95 % air with 5 % CO_2_ in Dulbecco’s modified essential medium (DMEM) (Gibco, Life Technologies) containing glucose (4.5 g/L), L-glutamine (2 mM), 1 mM sodium pyruvate and 50 µg/ml uridine, fetal bovine serum (FBS) (10% v/v), penicillin (100 U/ml), and streptomycin sulphate (100 µg/ml). The cells were tested and shown to be free off mycoplasma contamination. HEK 293T cells were plated at 60 % confluence in 15 cm plates and transfected with mammalian expression plasmids in OptiMEM media (Invitrogen). 158 ng/cm^2^ of NSUN4 and MTERF4 plasmid DNA, in equal ratios, were transfected using Fugene HD (Roche). Cell incubations were carried out for 72 h following transfection and mitochondria were isolated as described previously^3^.

### Sucrose gradients of mitochondrial ribosomes to analyze NSUN4 mutants

Sucrose gradient fractionation was carried out on purified mitochondria as previously described^3^.

### Immunoblotting

Specific proteins were detected using rabbit monoclonal antibodies against: MRPL12 (bL12m) (16394-1-AP), MRPS16 (bS16m) (16735-1-AP), and FLAG (Sigma, F7425); and mouse monoclonal antibodies against HA (Cell Signaling, 2367). All primary antibodies were diluted 1:1000 using the Odyssey blocking buffer (LI-COR). IR Dye 800CW Goat Anti-Rabbit IgG or IR Dye 680LT Goat Anti-Mouse IgG (LI-COR) secondary antibodies (diluted 1:10,000) were used and the immunoblots were visualized using the Odyssey Infrared Imaging System (LI-COR).

### Cryo-EM sample preparation and data acquisition

Quantifoil R2/2 holey carbon copper grids (Quantifoil Micro Tool) were prepared by applying an additional thin layer of continuous carbon, followed by glow-discharging for 15 s at 15 mA using an easiGlow Discharge cleaning system (PELCO). For both datasets, 4 µl of resuspended MTG1-3xFLAG tagged mitoribosome sample was directly applied onto the grid mounted in the Vitrobot chamber (Thermo Fisher Scientific) and incubated for 1 min. Excess of buffer was blotted away, and the grid was immediately plunge frozen in 1:2 ethane:propane (Carbagas) at liquid nitrogen temperature. The Vitrobot chamber was kept at 4 °C and 100 % humidity during the whole procedure. For the second dataset, NP-40 detergent was added to a final concentration of 0.001 % just before applying the MTG1-3xFLAG tagged mitoribosome sample onto the grid.

Both datasets were collected on a Titan Krios cryo-transmission electron microscope (Thermo Fisher Scientific) operating at 300 kV. For the first dataset, the microscope was equipped with a Falcon IIIEC Direct Electron Detector (FEI) and the movies were collected in integrating mode with a pixel size of 1.087 Å/pix, 30 frames and a total dose of 60 e^-^/Å^2^, with defocus varying from -3.0 µm to -0.6 µm. The second dataset was collected on a microscope equipped with a K3 detector (Gatan), mounted to a GIF Quantum LS imaging filter operated with an energy filter slit width of 20 eV. The movies were collected in counting and super-resolution mode, with 40 frames and a total dose of 60 e^-^/Å^2^ at a physical pixel size of 1.06 Å/pix (0.53 Å/pix in super resolution) with defocus varying from -3.0 µm to - 0.6 µm. Collection of both datasets was automated with the EPU software (Thermo Fisher Scientific).

### Cryo-EM data processing

Both datasets were collected and processed independently. Unless stated otherwise, all processing steps were performed using cryoSPARC 3.1^4^.

Movies of the first dataset collected on the Falcon IIIEC Direct Electron Detector (FEI) were drift-corrected and dose-weighted using MotionCor2^5^, and the corrected 15,441 micrographs were imported into cryoSPARC. The CTF parameters for each micrograph were estimated using Patch-Based CTF Estimation, and 100 randomly selected micrographs were used to pick initial particles using a Laplacian-of-Gaussian filter-based method. These particles were subjected to a 2D classification. Classes resembling the large subunit of the human mitoribosome were used as a reference for picking particles from the whole dataset. The resulting 2,214,072 particles were extracted at 5.6-fold binning and subjected to 2D classification. Good-looking classes were selected and used for an *ab-initio* reconstruction to create the initial 3D model. The obtained model was then used as an input in homogenous refinement with all 1,064,609 selected particles from the 2D classification. The resulting refined map was used to create a mask covering the intersubunit side of the large subunit in UCSF Chimera^6^. Aligned particles, together with the mask, were used in a 3D Variability analysis^7^ with 8 modes to solve and the resolution filtered to 6 Å. The particles were then divided into 10 clusters using 3D Variability Display.

The second dataset was pre-processed during collection using cryoSPARC live^4^. The pre-processing included 2-fold binning, drift-correction, dose-weighting, CTF estimation and particle picking using a Laplacian-of-Gaussian filter-based method, resulting in 10,682 processed micrographs and 1,626,837 extracted particles that were exported into cryoSPARC. The particles were then subjected to 2D classification, and classes resembling the mitoribosomal large subunit were selected, resulting in 833,994 particles. All particles were used for *ab-initio* reconstruction, and the resulting cryo-EM map was used as an input model in homogenous refinement. The previously created intersubunit mask was resampled onto the new map using UCSF Chimera^6^ and together with the aligned particles was used for 3D Variability analysis^7^ with 5 modes to solve and the resolution filtered to 10 Å. The particles were then divided into 25 clusters using 3D Variability Display.

Overall, processing of both datasets resulted in similarly looking 3D classes. The only difference were states B and D0’’, which were present only in the first and the second dataset, respectively. The key classes found in both datasets were re-extracted at full-size and refined again, this time with per-particle defocus estimation^8^. The local resolution was calculated using a locally windowed FSC method as described in^9^.

### Model building and refinement

Published structures of the mature human mitoribosome^10^ (PDB 6ZM6) and of a late assembly intermediate containing the ACP module^11^ (PDB 5OOL) were used as initial models and docked into the cryo-EM maps using UCSF Chimera^6^. Composite models were assembled in PyMOL Molecular Graphics System, Version 2.1.5 (Schrödinger, LLC), followed by manual rebuilding of the proteins and nucleic acids using Coot^12^. For interpretation of the additional density features representing the assembly factors, the crystal structures of the human NSUN4–MTERF4 dimer^13^ (PDB 4FZV) and MRM2 (PDB 2NYU) were fitted and readjusted. For MTG2, an initial model was obtained using the Phyre2 modelling server^14^ based on a crystal structure of the *E. coli* homolog ObgE^15^ (PDB 5M04). The N-terminal Obg and C-terminal G domains were docked individually and rebuilt. For fitting of the GDP and SAM cofactors, superimposed high-resolution structures were used as a guide.

The models were real space refined for 5 cycles using Phenix version 1.19.1^16^ while applying side chain rotamer and Ramachandran restraints. Remaining discrepancies between the models and maps were detected and corrected using real space difference density maps and the geometry validation tools implemented in Coot^12^. The final model geometry was validated using MolProbity^17^ (**Supplementary Table 1**). To evaluate the quality of the fit of the refined models to the EM maps, real space correlation coefficients (CC_mask_) as well as the model versus map FSCs at the FSC = 0.5 criterion were calculated. The resulting resolutions were close to those calculated from the map half-sets at the FSC = 0.143 criterion (**Supplementary Fig. 3**).

## Data availability

The atomic coordinates were deposited in the RCSB Protein Data Bank (PDB) under accession numbers xxxx, xxxx and xxxx. The cryo-EM maps were deposited in the Electron Microscopy Data Bank (EMDB) under accession numbers EMD-xxxx, EMD-xxxx, EMD-xxxx, EMD-xxxx, EMD-xxxx and EMD-xxxx.

## Figures

Molecular graphics and analyses were performed with UCSF Chimera^6^ and UCSF ChimeraX^18^, developed by the Resource for Biocomputing, Visualization, and Informatics at the University of California, San Francisco, with support from National Institutes of Health P41-GM103311 and R01-GM129325 and the Office of Cyber Infrastructure and Computational Biology, National Institute of Allergy and Infectious Diseases. Detailed views of cryo-EM map densities were created using The PyMOL Molecular Graphics System, Version 2.1.5 Schrödinger, LLC.

## Acknowledgments

Authors thank A. Jomaa, E. Kummer and S. Mattei for helpful discussions and creating a great working atmosphere. T.L. thanks R. Jost for help with the sample preparation, A. Picenoni and E. Kummer for help with visualizing the sample using negative stain EM, and A. Ries for establishing the cell culture laboratory. All authors would like to thank the ETH Zürich scientific centre for optical and electron microscopy (ScopeM) for access to electron microscopy equipment.

## Authors’ contributions

N.B. and T.L. designed the project. T.L., T.S. and M.J. prepared the sample for cryo-EM. T.S. and M.S. established the methods for transfection of HEK cells with affinity tagged mitochondrial proteins. A.S. and T.L. collected the cryo-EM data. M.J. and T.L. processed the cryo-EM data. M.L. built and performed coordinate refinement of the atomic models. T.L. and M.J. interpreted the structures. O.R., R.L. and A.F. performed *in vivo* experiments in HEK cells. T.L., M.J., A.F. and N.B. drafted the initial manuscript, all authors contributed to the final version.

## References

1. Kummer, E. & Ban, N. Mechanisms and regulation of protein synthesis in mitochondria. Nat. Rev. Mol. Cell Biol., (2021) doi.org/10.1038/s41580-021-00332-2.

2. Itoh, Y. et al. Mechanism of membrane-tethered mitochondrial protein synthesis. Science 371, 846–849 (2021).

3. Greber, B. J. et al. The complete structure of the 55S mammalian mitochondrial ribosome. Science 348, 303–308 (2015).

4. Amunts, A., Brown, A., Toots, J., Scheres, S. H. W. & Ramakrishnan, V. The structure of the human mitochondrial ribosome. Science 348, 95–98 (2015).

5. Bieri, P., Greber, B. J. & Ban, N. High-resolution structures of mitochondrial ribosomes and their functional implications. Curr. Opin. Struct. Biol. 49, 44–53 (2018).

6. Ferrari, A., Del’Olio, S. & Barrientos, A. The Diseased Mitoribosome. FEBS Lett. (2020) doi:10.1002/1873-3468.14024.

7. Bogenhagen, D. F., Ostermeyer-Fay, A. G., Haley, J. D. & Garcia-Diaz, M. Kinetics and Mechanism of Mammalian Mitochondrial Ribosome Assembly. Cell Rep. 22, 1935–1944 (2018).

8. Maiti, P., Lavdovskaia, E., Barrientos, A. & Richter-Dennerlein, R. Role of GTPases in Driving Mitoribosome Assembly. Trends Cell Biol. 1–14 (2021) doi:10.1016/j.tcb.2020.12.008.

9. Silva, D. De, Tu, Y.-T., Amunts, A., Fontanesi, F. & Barrientos, A. Mitochondrial ribosome assembly in health and disease. Cell Cycle 14, 2226–2250 (2015).

10. Brown, A. et al. Structures of the human mitochondrial ribosome in native states of assembly. Nat. Struct. Mol. Biol. 24, 866–869 (2017).

11. Kotani, T., Akabane, S., Takeyasu, K., Ueda, T. & Takeuchi, N. Human G-proteins, ObgH1 and Mtg1, associate with the large mitochondrial ribosome subunit and are involved in translation and assembly of respiratory complexes. Nucleic Acids Res. 41, 3713–3722 (2013).

12. Kim, H.-J. & Barrientos, A. MTG1 couples mitoribosome large subunit assembly with intersubunit bridge formation. Nucleic Acids Res. 46, 8435–8453 (2018).

13. Cámara, Y. et al. MTERF4 Regulates Translation by Targeting the Methyltransferase NSUN4 to the Mammalian Mitochondrial Ribosome. Cell Metab. 13, 527–539 (2011).

14. Metodiev, M. D. et al. NSUN4 Is a Dual Function Mitochondrial Protein Required for Both Methylation of 12S rRNA and Coordination of Mitoribosomal Assembly. PLoS Genet. 10, e1004110 (2014).

15. Yakubovskaya, E. et al. Structure of the Essential MTERF4:NSUN4 Protein Complex Reveals How an MTERF Protein Collaborates to Facilitate rRNA Modification. Structure 20, 1940–1947 (2012).

16. Spåhr, H., Habermann, B., Gustafsson, C. M., Larsson, N.-G. & Hallberg, B. M. Structure of the human MTERF4–NSUN4 protein complex that regulates mitochondrial ribosome biogenesis. Proc. Natl. Acad. Sci. 109, 15253–15258 (2012).

17. Rorbach, J. et al. MRM2 and MRM3 are involved in biogenesis of the large subunit of the mitochondrial ribosome. Mol. Biol. Cell 25, 2542–2555 (2014).

18. Lee, K.-W. & Bogenhagen, D. F. Assignment of 2′-O-Methyltransferases to Modification Sites on the Mammalian Mitochondrial Large Subunit 16S Ribosomal RNA (rRNA). J. Biol. Chem. 289, 24936–24942 (2014).

19. Nikolay, R. et al. Snapshots of native pre-50S ribosomes reveal a biogenesis factor network and evolutionary specialization. Mol. Cell 81, 1200–1215.e9 (2021).

20. Maiti, P., Antonicka, H., Gingras, A. C., Shoubridge, E. A. & Barrientos, A. Human GTPBP5 (MTG2) fuels mitoribosome large subunit maturation by facilitating 16S rRNA methylation. Nucleic Acids Res. 48, 7924–7943 (2020).

21. Kummer, E. et al. Unique features of mammalian mitochondrial translation initiation revealed by cryo-EM. Nature 560, 263–267 (2018).

22. Jaskolowski, M. et al. Structural Insights into the Mechanism of Mitoribosomal Large Subunit Biogenesis. Mol. Cell 79, 629–644.e4 (2020).

23. Soufari, H. et al. Structure of the mature kinetoplastids mitoribosome and insights into its large subunit biogenesis. Proc. Natl. Acad. Sci. U. S. A. 117, 29851–29861 (2020).

24. Tobiasson, V. et al. Interconnected assembly factors regulate the biogenesis of mitoribosomal large subunit. EMBO J. (2021) doi:10.15252/embj.2020106292.

25. Saurer, M. et al. Mitoribosomal small subunit biogenesis in trypanosomes involves an extensive assembly machinery. Science. 365, 1144–1149 (2019).

26. Sun, N., Youle, R. J. & Finkel, T. The Mitochondrial Basis of Aging. Mol. Cell 61, 654–666 (2016).

27. Pearce, S., Nezich, C. L. & Spinazzola, A. Mitochondrial diseases: Translation matters. Mol. Cell. Neurosci. 55, 1–12 (2013).

28. Boczonadi, V. & Horvath, R. Mitochondria: Impaired mitochondrial translation in human disease. Int. J. Biochem. & Cell Biol. 48, 77–84 (2014).

## Methods references

1. Clayton, D. A. & Shadel, G. S. Isolation of Mitochondria from Cells and Tissues. Cold Spring Harb. Protoc. 2014, pdb.top074542--pdb.top074542 (2014).

2. Aibara, S., Andréll, J., Singh, V. & Amunts, A. Rapid Isolation of the Mitoribosome from HEK Cells. J. Vis. Exp. (2018) doi:10.3791/57877.

3. Lee, R. G. et al. Cardiolipin is required for membrane docking of mitochondrial ribosomes and protein synthesis. J. Cell Sci. 133, jcs240374 (2020).

4. Punjani, A., Rubinstein, J. L., Fleet, D. J. & Brubaker, M. A. cryoSPARC: algorithms for rapid unsupervised cryo-EM structure determination. Nat. Methods 14, 290–296 (2017).

5. Zheng, S. Q. et al. MotionCor2: Anisotropic correction of beam-induced motion for improved cryo-electron microscopy. Nat. Methods 14, 331–332 (2017).

6. Pettersen, E. F. et al. UCSF Chimera - A visualization system for exploratory research and analysis. J. Comput. Chem. 25, 1605–1612 (2004).

7. Punjani, A. & Fleet, D. J. 3D variability analysis: Resolving continuous flexibility and discrete heterogeneity from single particle cryo-EM. J. Struct. Biol. 213, 107702 (2021).

8. Zivanov, J., Nakane, T., Scheres, S. H. W. & Em, C. Estimation of high-order aberrations and anisotropic magnification from cryo-EM data sets in RELION-3.1. IUCrJ 7, 253–267 (2020).

9. Cardone, G., Heymann, J. B. & Steven, A. C. One number does not fit all: Mapping local variations in resolution in cryo-EM reconstructions. J. Struct. Biol. 184, 226–236 (2013).

10. Itoh, Y. et al. Mechanism of membrane-tethered mitochondrial protein synthesis. Science 371, 846–849 (2021).

11. Brown, A. et al. Structures of the human mitochondrial ribosome in native states of assembly. Nat. Struct. Mol. Biol. 24, 866–869 (2017).

12. Emsley, P., Lohkamp, B., Scott, W. G. & Cowtan, K. Features and development of Coot. Acta Crystallogr. Sect. D Biol. Crystallogr. 66, 486–501 (2010).

13. Yakubovskaya, E. et al. Structure of the Essential MTERF4:NSUN4 Protein Complex Reveals How an MTERF Protein Collaborates to Facilitate rRNA Modification. Structure 20, 1940–1947 (2012).

14. Kelley, L. A., Mezulis, S., Yates, C. M., Wass, M. N. & Sternberg, M. J. E. E. The Phyre2 web portal for protein modeling, prediction and analysis. Nat. Protoc. 10, 845–858 (2015).

15. Gkekas, S. et al. Structural and biochemical analysis of Escherichia coli ObgE, a central regulator of bacterial persistence. J. Biol. Chem. 292, 5871–5883 (2017).

16. Afonine, P. V. et al. Real-space refinement in PHENIX for cryo-EM and crystallography. Acta Crystallogr. Sect. D, Struct. Biol. 74, 531–544 (2018).

17. Williams, C. J. et al. MolProbity: More and better reference data for improved all-atom structure validation. Protein Sci. 27, 293–315 (2018).

18. Goddard, T. D. et al. UCSF ChimeraX: Meeting modern challenges in visualization and analysis. Protein Sci. 27, 14–25 (2018).

