## Supplementary Info for "Stepwise maturation of the peptidyl transferase region of human mitoribosomes"

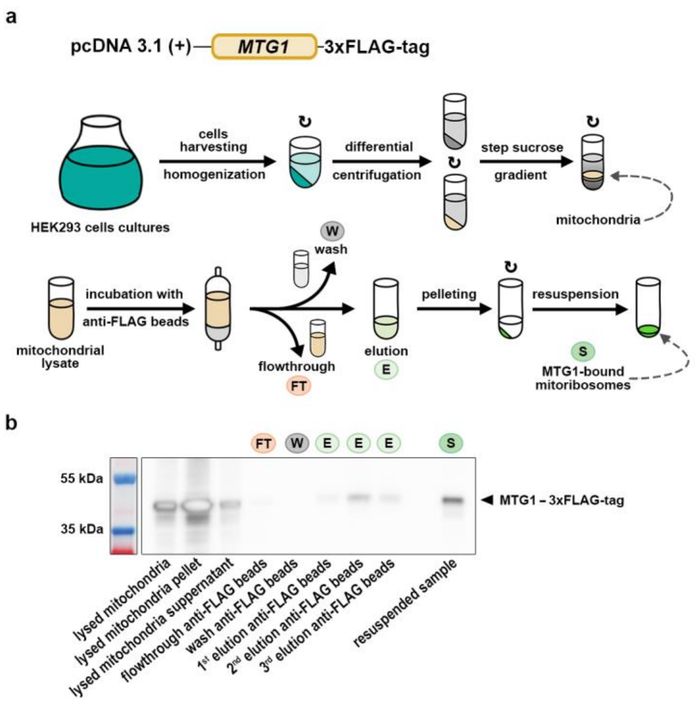


**Supplementary Figure 1. 3x-FLAG tagged MTG1 co-pellets with the mitoribosomal particles.** (a) Schematic representation of the mitochondria preparation, followed by the purification of mitoribosomes *via* C-terminally 3x-FLAG-tagged MTG1**.** Curved arrows indicate an (ultra)centrifugation step. (b) Western Blot analysis of the purification procedure using antibodies against the 3xFLAG tag.

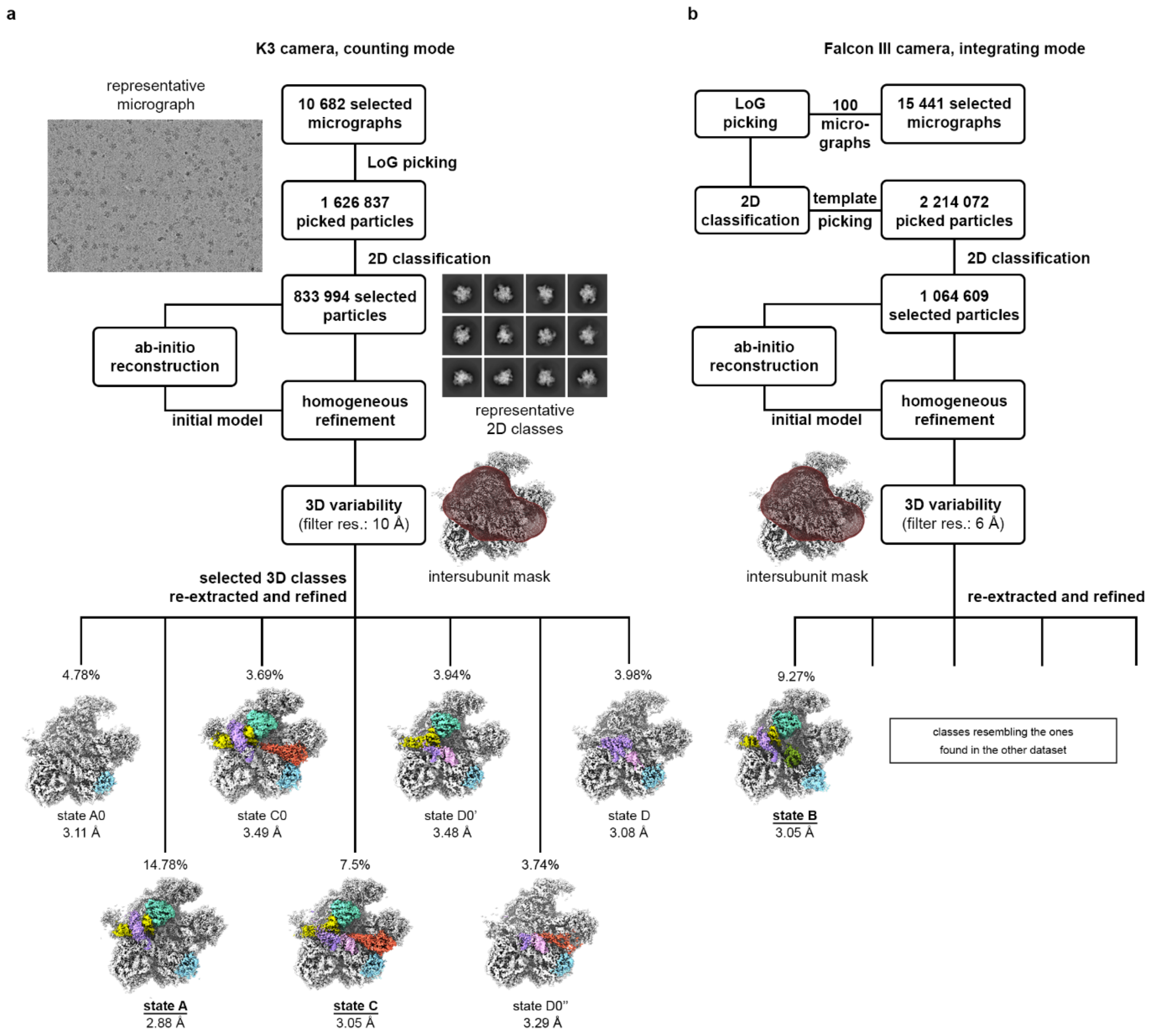


**Supplementary Figure 2. Data processing of the purified particles bound to the tagged MTG1 assembly factor.** Data processing schemes for both datasets collected from the particles obtained by affinity purification of C-terminally tagged MTG1 assembly factor. Both datasets, collected using K3 (a) and Falcon IIIEC (b) cameras, were processed independently. Cryo-EM maps of selected 3D classes with density of assembly factors as well as key rRNA regions color-coded as in Fig. 1 are shown.

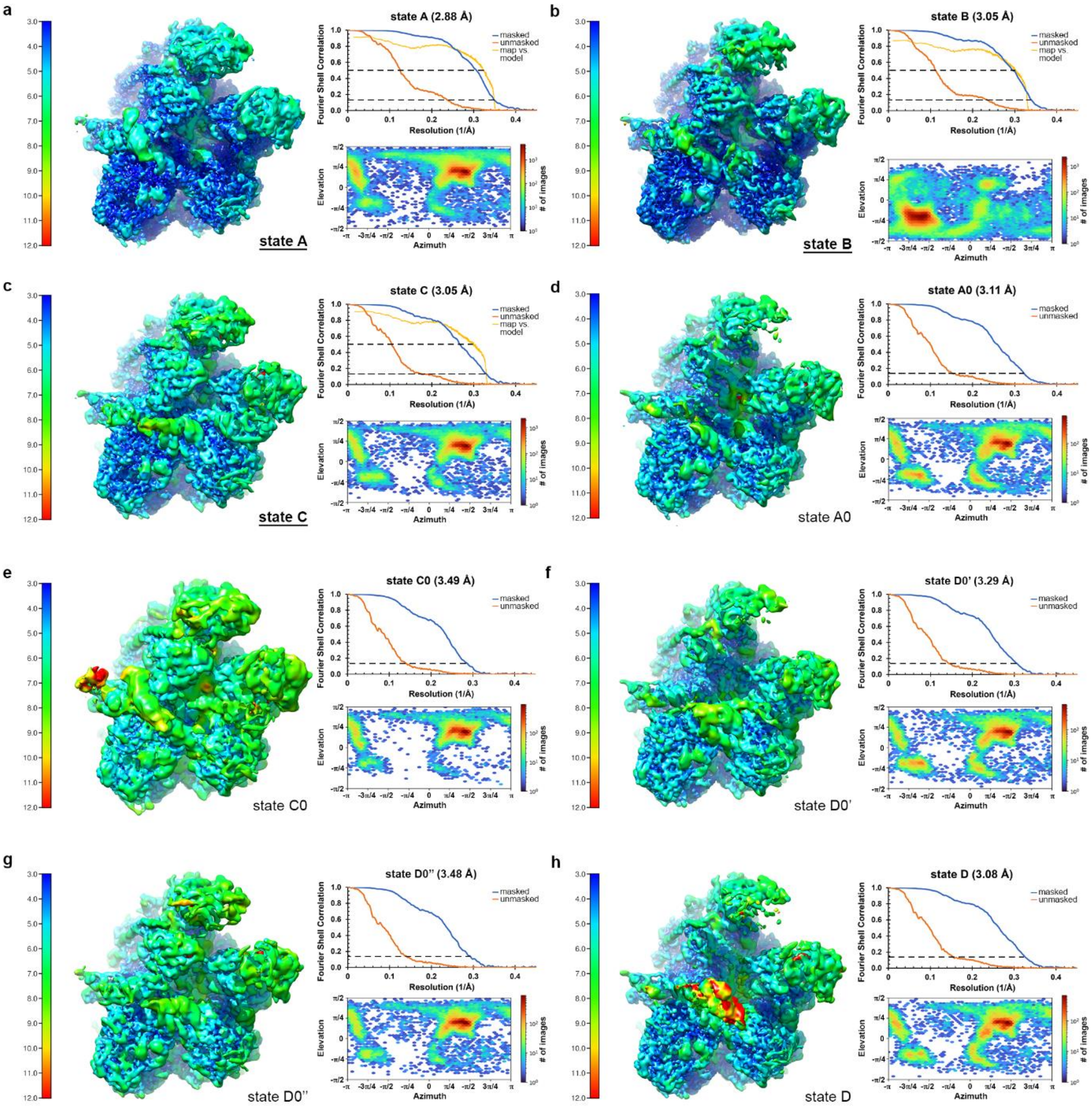


**Supplementary Figure 3. Cryo-EM data statistics.** Cryo-EM maps, colored according to the local resolution with scale bars presented on the left of each panel, are shown for state A (a), state B (b), state C (c), state A0 (d), state C0 (e), state D0' (f), state D0'' (g) and state D (h). Each panel contains the FSC curve on the top right and the particles angular distribution graph on the bottom right.

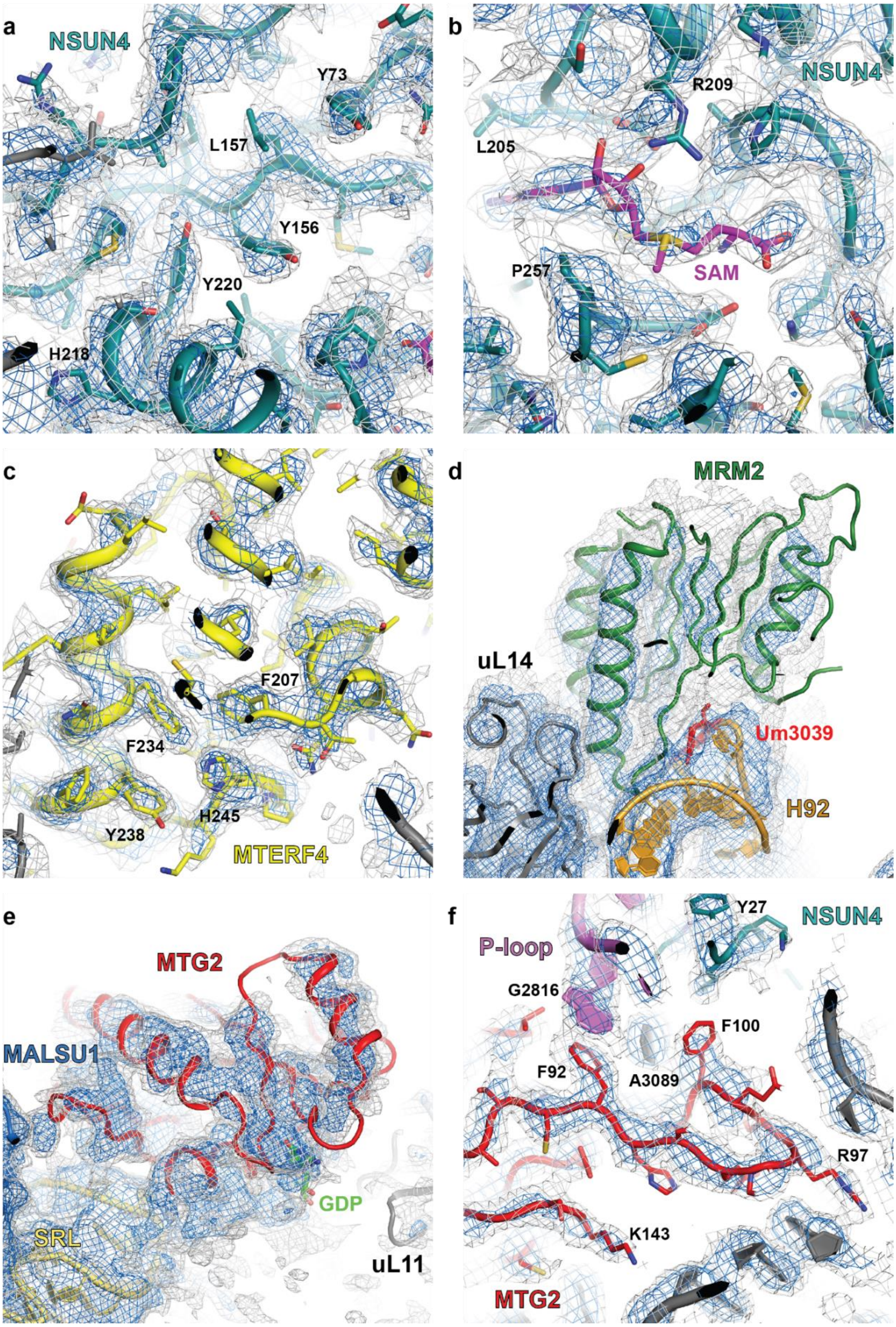


**Supplementary Figure 4. Examples of electron density maps in the areas of the bound maturation factors.** Overview of the EM density for (a) NSUN4, (b) the NSUN4 S-adenosyl-methionine cofactor, (c) MTERF4, (d) MRM2, (e) the G domain of MTG2 and (f) the N-terminal Obg domain of MTG2 and NSUN4 interacting with the P loop.

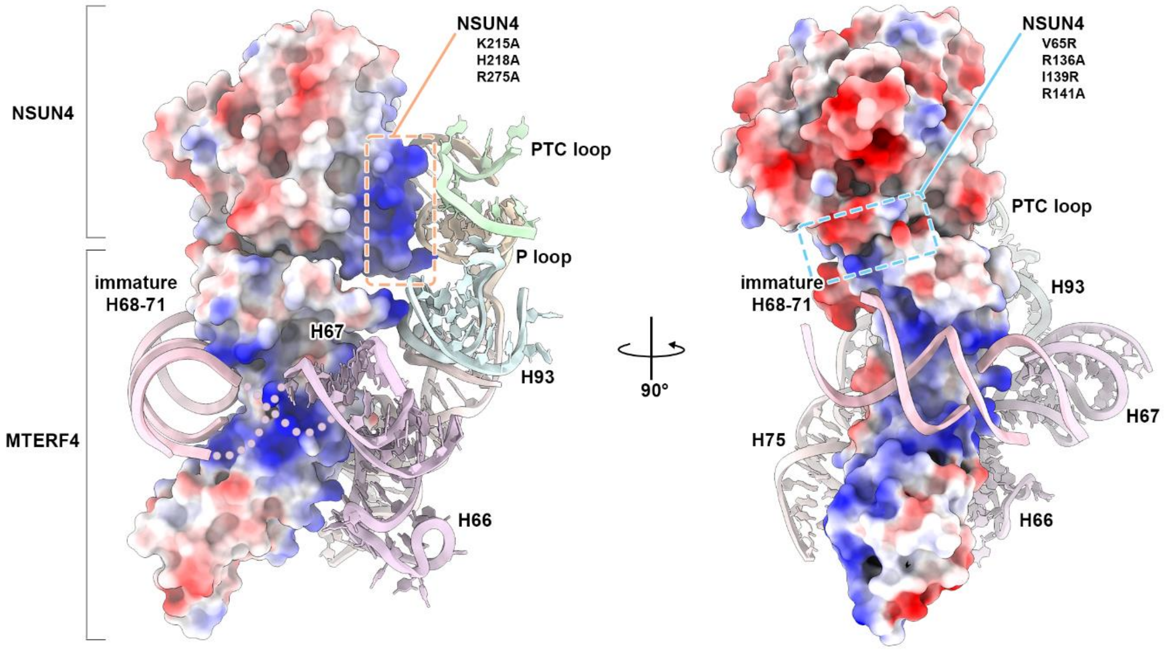


**Supplementary Figure 5. NSUN4 binding to key elements of the PTC rRNA region and MTERF4 stabilizing a distinct immature conformation of H68-71.** Atomic models of NSUN4–MTERF4 are shown in surface representation and colored according to their electrostatic potential. Selected rRNA elements that interact with NSUN4–MTERF4 are shown in cartoon and are labeled individually. NSUN4–MTERF4 is shown as viewed from the GTPase associated center (left) and from intersubunit space (right). Regions of introduced NSUN4 mutations are indicated with dashed line boxes.

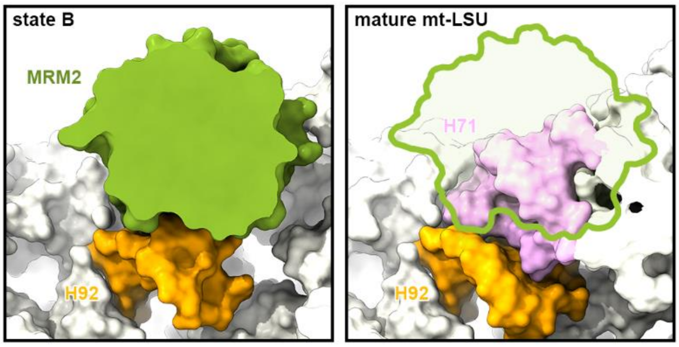


**Supplementary Figure 6. MRM2 methylates U3039 in the A-loop before maturation of H71.** Cross-sections of the atomic models of state B (left) and the mature mt-LSU (right, PDB: 6ZM6) are shown as viewed from the CP. The MRM2 methyltransferase, H92 (A loop) and H71 are labeled and colored individually. On the right panel, the green outline represents superposed MRM2 to visualize steric hindrance of the assembly factor with the mature H71.

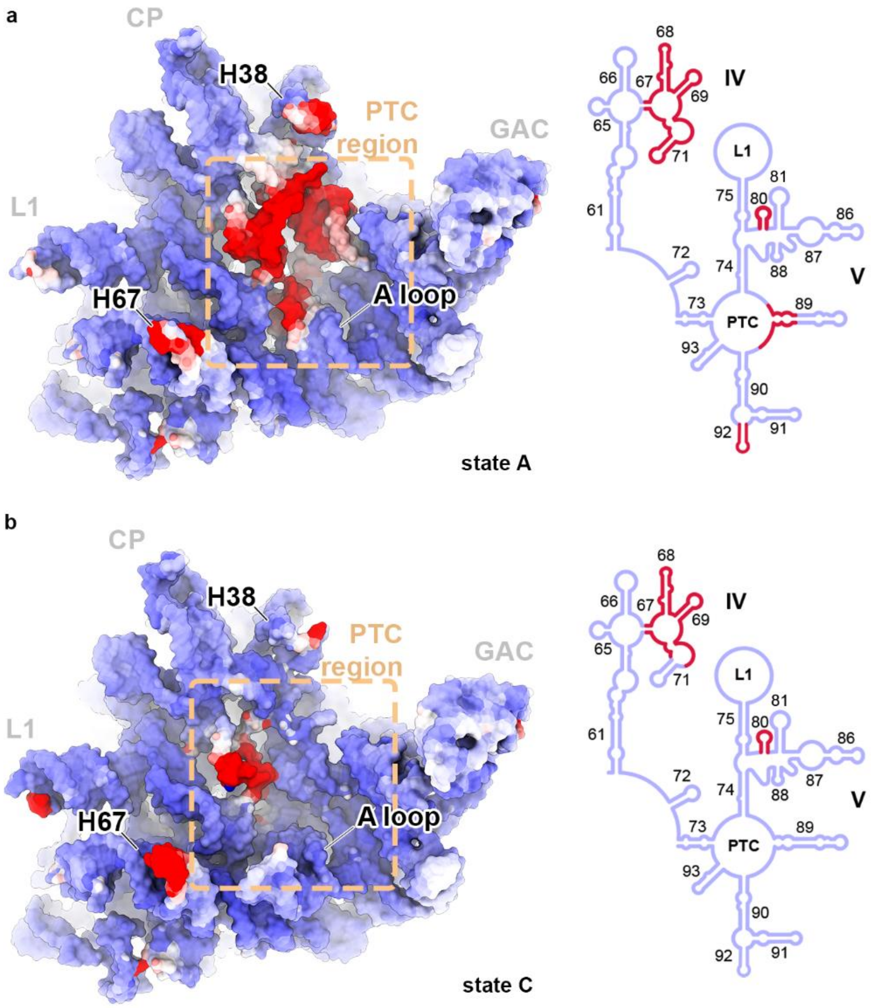


**Supplementary Figure 7. Stepwise maturation of the key rRNA regions of the mt-LSU.** The 16S rRNA atomic models of states A (a) and C (b) are shown in surface representation and colored according to the root-mean-square deviation (RMSD) compared to the mature 16S rRNA (PDB: 6ZM6). RMSD values range from 0 Å (blue) to 8 Å and above (red). Corresponding 2D diagrams of 16S rRNA domains IV and V are shown on the right of each panel and colored according to their respective atomic model.

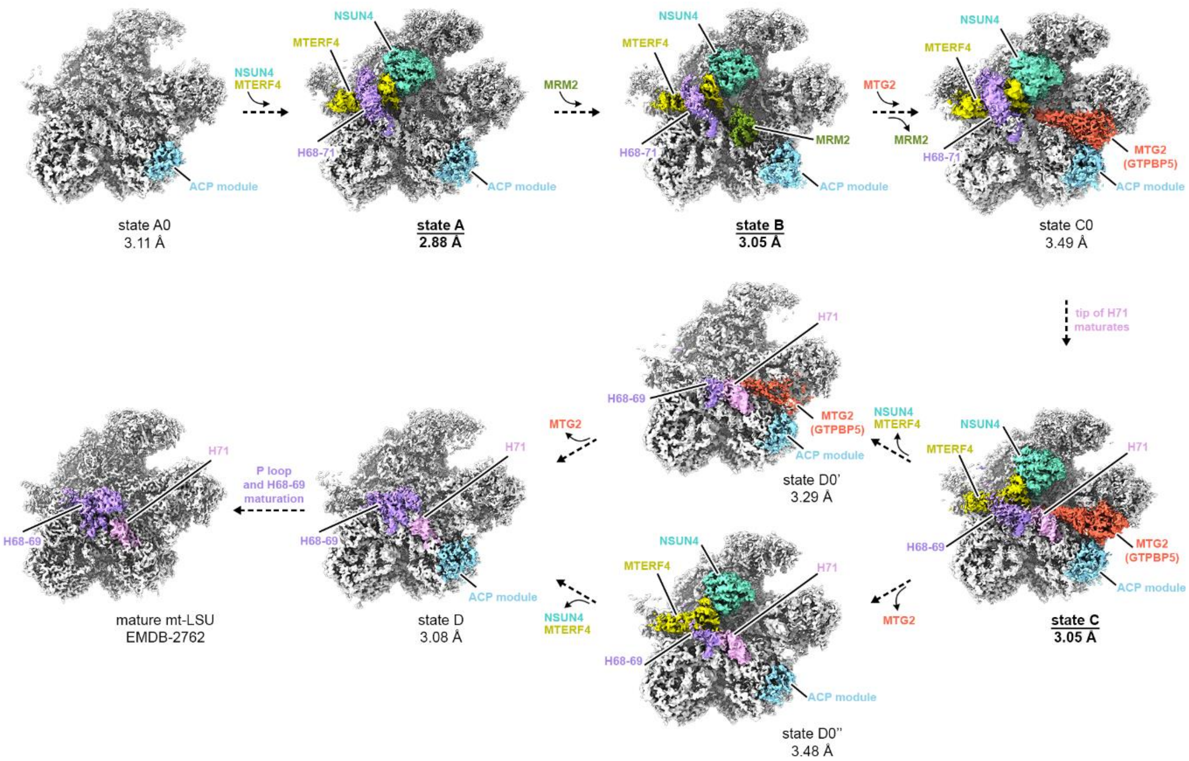


**Supplementary Figure 8. Stepwise maturation of the mt-LSU including all obtained cryo-EM maps.** The pathway of mt-LSU maturation is proposed based on all obtained cryo-EM maps. The densities corresponding to the assembly factors and important rRNA fragments are colored and labeled individually. The ACP module includes MALSU1, L0R8F8 and mt-ACP.

**Supplementary Table 1. Cryo-EM data collection, refinement and validation statistics.**

| | **State A MTERF-NSUN4 + ACP module** | **State B MTERF-NSUN4-MRM2 + ACP module** | **State C MTERF-NSUN4-MTG2-H71 + ACP module** |
|---|---|---|---|
| EMDB code | #### | #### | #### |
| PDB code | #### | #### | #### |
| **Data collection and processing** | | | |
| Camera | K3 | Falcon IIIEC | K3 |
| Magnification | 81,000 (nominal) | 75,000 (nominal) | 81,000 (nominal) |
| Voltage (kV) | 300 | | |
| Electron exposure ($e^{-}$/Å$^2$) | 60 | | |
| Defocus range (μm) | 0.6-3.0 | | |
| Pixel size (Å) | 1.06 (super-res. pix. at 0.53Å/pix.) | 1.087 | 1.06 (super-res. pix. at 0.53Å/pix.) |
| Initial particle images (no.) | 833,994 | 1,064,609 | 833,994 |
| Final particle images (no.) | 123,285 | 114,557 | 62,565 |
| Map resolution at FSC=0.143 (Å) | 2.9 | 3.1 | 3.1 |
| **Structure refinement in PHENIX 1.19.1** | | | |
| Model resolution at FSC=0.5 (Å) | 3.1 | 3.3 | 3.4 |
| $CC_{mask}$ | 0.80 | 0.76 | 0.79 |
| Map sharpening B factor (Å$^2$) | – 64.1 | – 97.9 | – 54.6 |
| Model composition | | | |
| Non-hydrogen atoms | 107,683 | 108,570 | 110,405 |
| Protein residues | 9,274 | 9,374 | 9,588 |
| RNA residues | 1,516 | 1,516 | 1,521 |
| Ligands: $Mg^{2+}$ / $K^{+}$ / $Zn^{2+}$ / [$Fe_2$-$S_2$] / C10-PPT / SAM / GDP | 97 / 2 / 2 / 1 / 1 / 1 / - | 68 / 0 / 2 / 1 / 1 / 1 / - | 99 / 4 / 2 / 1 / 1 / 1 / 1 |
| B factors min/max/mean (Å$^2$) | | | |
| Protein | 14/122/57 | 20/246/91 | 27/356/101 |
| RNA | 8/183/52 | 22/382/77 | 27/278/72 |
| Ligand | 12/108/52 | 19/177/88 | 26/234/94 |
| R.m.s. deviations | | | |
| Bond lengths (Å) | 0.002 | 0.002 | 0.002 |
| Bond angles (°) | 0.408 | 0.402 | 0.403 |
| **Validation** | | | |
| MolProbity score | 1.3 | 1.3 | 1.3 |
| Clashscore | 5.1 | 5.3 | 5.3 |
| Poor rotamers (%) | 0.8 | 0.7 | 0.8 |
| Protein | | | |
| EM Ringer score | 3.2 | 2.5 | 2.8 |
| Ramachandran plot | | | |
| Favored (%) | 98.73 | 98.70 | 98.78 |
| Allowed (%) | 1.27 | 1.29 | 1.22 |
| Disallowed (%) | 0.0 | 0.01 | 0.0 |
| RNA | | | |
| Pucker outliers (%) | 0.5 | 0.7 | 0.6 |
| Bond outliers (%) | 0.0 | 0.0 | 0.0 |
| Angle outliers (%) | 0.0 | 0.0 | 0.0 |
| Suite outliers (%) | 17.1 | 18.4 | 18.5 |
